# Lack of Evidence for Local Adaptation in Risk Tolerance: A Genetic Study in Northern Senegal

**DOI:** 10.64898/2026.09.11.751078

**Authors:** Clément Mettling, Aby Mbengue, Catherine Breton, Gwen-Jiro Clochard, Birane Diouf, Charlotte Faurie, Omar Sene, Adline Delcamp, Erwan Guichoux, Guillaume Hollard, Marc Willinger, Michel Raymond

## Abstract

Risk tolerance is influenced by both environmental and genetic factors and may be shaped by local selective pressures in hazardous environments. In Northern Senegal, fishermen from the village of Guet Ndar are chronically exposed to high occupational mortality risk, making this population a potential model for local adaptation to risky environments. In a previous study, we showed that men from this fishing community were less risk-tolerant than men from a nearby farming village, although this difference could not be explained by variation at the DRD4 locus. Here, we investigated whether genetic variants previously associated with general risk tolerance in large genome-wide association studies (GWAS) exhibit signals of local adaptation in these Senegalese populations. We genotyped 44 candidate SNPs in 373 individuals sampled from the risky fishing area and the non-risky farming area. Population genetic analyses revealed extremely low and non-significant differentiation between the two populations (mean FST = 0.0003, p = 0.33), indicating that the two groups are genetically indistinguishable at these loci. Furthermore, none of the candidate SNPs showed a significant association with experimentally measured risk tolerance after correction for multiple testing. These results provide no evidence that local adaptation for risky behavior has occurred, several explanations may account for this negative result.

## Introduction

Risk tolerance varies substantially across individuals. While most people are risk-averse, others are more willing to engage in risky activities ^1^. This variation is partly shaped by cultural factors and may differ across populations and nationalities ^2^. Risk tolerance also varies with individual characteristics such as gender, age, and cognitive ability: women are generally less risk tolerant than men ^3–5^, individuals with higher cognitive ability tend to exhibit greater risk tolerance, and older individuals are typically less risk tolerant than younger ones ^6^. In addition to these environmental and demographic influences, risk tolerance has a genetic component. A large genome-wide association study (GWAS) identified 99 SNPs associated with general risk tolerance^7^. Consistent with this finding, previous studies have shown that risk-taking behavior reflects both phenotypic plasticity as individuals may adjust their risk preferences across contexts, and genetic influences ^8–12^.

The impact of background risk on risk-taking is theoretically ambiguous. Under Expected Utility theory, an unavoidable and uninsurable background risk generally increases aversion to independent risks, implying that independent risks act as substitutes: individuals exposed to a hazardous environment become less willing to take additional risks ^13^. By contrast, several non-Expected Utility models predict weaker, ambiguous, or even opposite effects, in which case independent risks may act as complements, so that exposure to one source of risk increases willingness to engage in another ^14^. Empirical evidence nevertheless suggests that exposure to background risk often reduces risk-taking behavior ^11,15,16^. Because risk tolerance is partly heritable, persistent exposure to hazardous environments may favor individuals with particular risk-related behavioral traits, potentially leading to local genetic adaptation.

In a previous study ^17^, we compared men from the fishing community of Guet Ndar (labeled the risky area below), located in the Saint-Louis region of northern Senegal, with men from a nearby farming village (non-risky area). In Guet Ndar, fishing, a highly hazardous activity associated with approximately 20 fatalities per year, constitutes the primary occupation of about 80% of the adult male workforce and was therefore designated as the risky area, whereas the farming village served as the non-risky area. Risk tolerance was measured using an incentivized lottery task ^18^, in which participants could win money by choosing between cards corresponding to lotteries characterized by an increasing average amount won but also an increasing variance.

We found that individuals from the risky area were significantly less risk tolerant than those from the non-risky area. We also observed a genetic association at the dopamine receptor DRD4 locus, with carriers of the 7R allele displaying higher risk tolerance. However, this genetic effect could not account for the behavioral differences between the two areas because the frequency of the 7R allele was similar in both populations^17^.

Given the persistent exposure to occupational risk in Guet Ndar, this population provides a potentially informative setting for studying local adaptation. In particular, the village appears to satisfy several conditions that may favor genetic adaptation: limited migration, substantial economic benefits associated with engaging in a risky occupation, and the existence of heritable variation in risk-related traits ^19^. Local adaptation has already enabled in a small population the isolation of SNPs linked to height and BMI, complex traits influenced by hundreds of genes, each with a modest effect that typically requires hundreds of thousands of individuals for detection^20^. We therefore tested whether genetic variants previously associated with general risk tolerance in a large GWAS ^7^ show evidence of local adaptation in this population.

## Results and Discussion

We selected 83 SNPs from the GWAS study based on technical criteria for MassArray genotyping, using a total of 380 samples (190 from each area). We excluded those SNP which were monomorphic (12 SNP), with ambiguous signals (12 SNP) or with low coverage (<80 %, 15 SNP) and analyzed the 44 remaining SNP for the genotypic differentiation between the two areas. We removed 7 samples with low DNA recovery showing less than 11 positive genotyping out of 44 (i.e. N=186 of the risky area and N=187 in the non-risky area). Hardy-Weinberg equilibrium was rejected in each area, with a heterozygote deficiency (Fis = 0.12). A PCA plot of genetic variation for 44 SNPs shows no evidence of population stratification (Fig. 1).

**Fig. 1:**
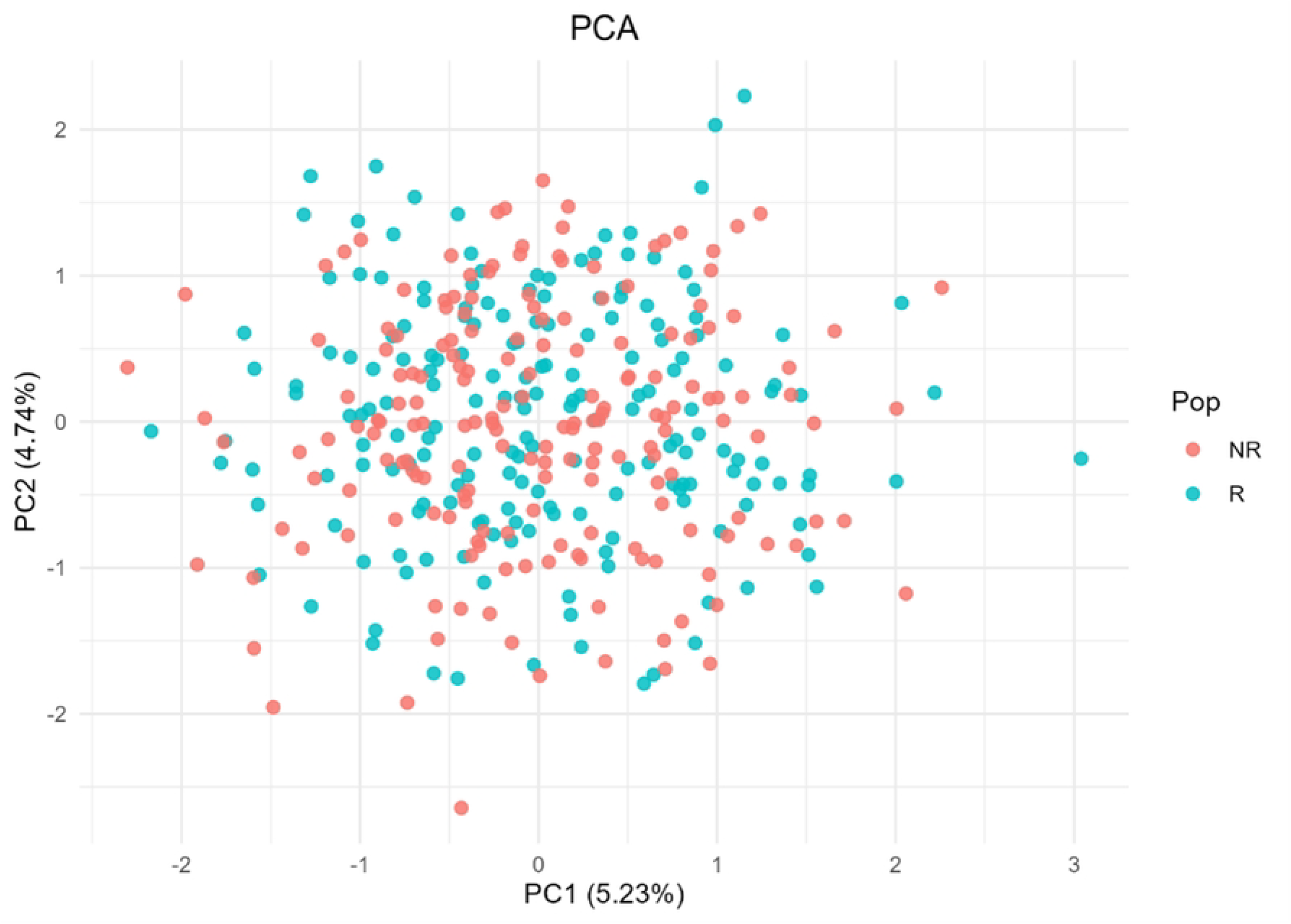
PCA Plot of Genetic Variation for 44 SNPs in Risky(R) vs Non-Risky (NR) area.

The genotypic differentiation between the two areas was measured as a mean Fst of 0.0003 and is non-significant with a p-value of 0.33. The distribution of the 44 SNP Fst is shown in Fig. 2.

**Fig. 2:**
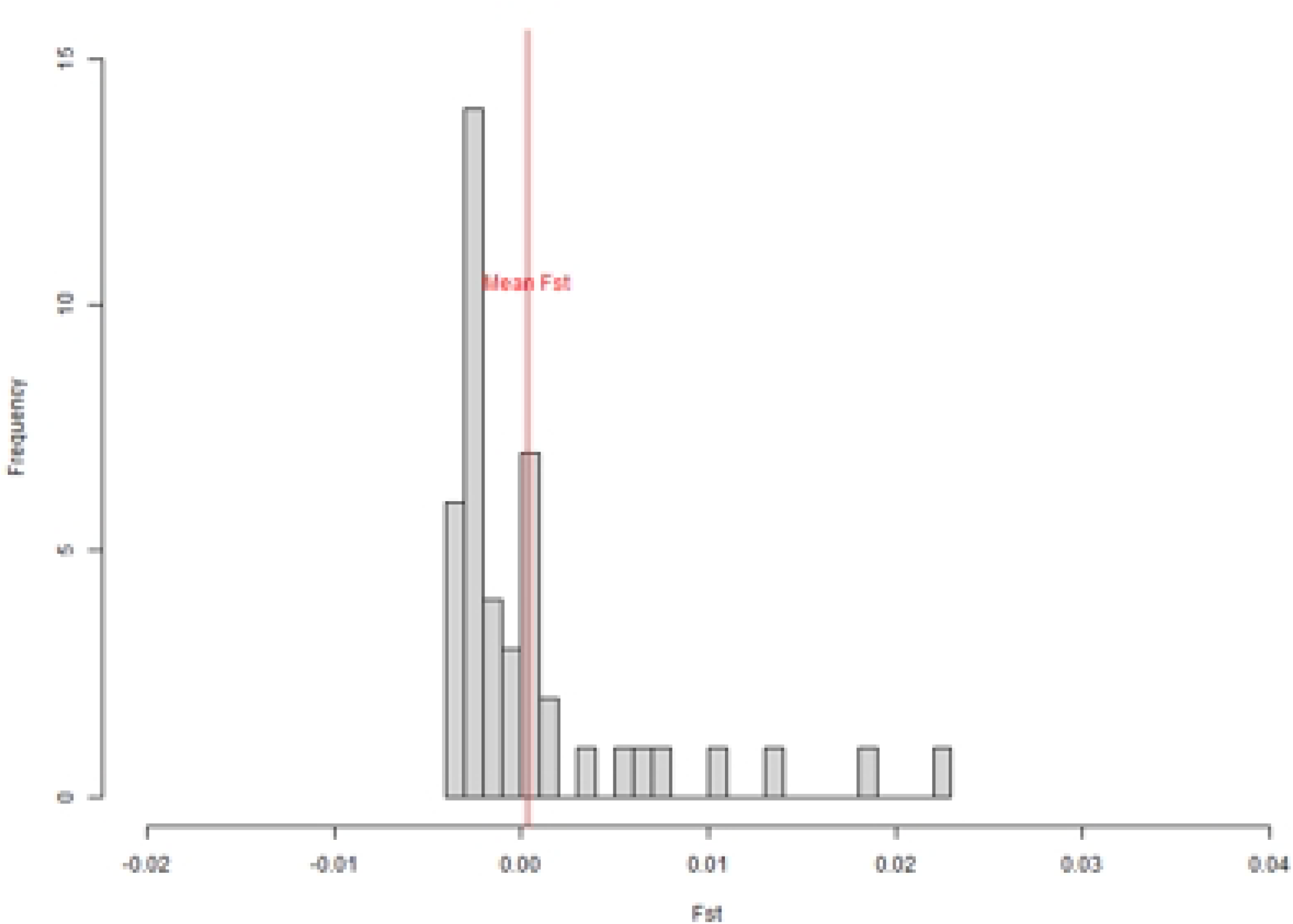
Fst distribution between the 2 areas across the 44 SNPs.

None of the SNPs showed a significant differentiation after correcting for multiple testing (Table S1). This level of genotypic differentiation was similar with those displayed by the 30 microsatellites analyzed in our previous study and confirms that the populations from the two villages are genetically indistinguishable.

We then tested the additive and dominance contribution of each SNP to risk tolerance (coded as 0 for maximum risk to 1 for no risk) in a Generalized Linear Model (glm). We adjusted for potentially confounding variables: area, education, income and age (Table S2). After correcting for multiple hypothesis testing, none of the SNPs showed a significant genetic contribution to risk tolerance. Fig. 3 shows the distribution of the - log of the p-value for additive and dominant contribution for each SNP, each one is below the line showing Bonferroni correction. A censored model (Tobit) gives analogous results (data not shown). Of the four adjustment variables, only area consistently had p-values below 0.012 (though still above the Bonferroni threshold) across all 44 SNPs (Table S2), reflecting differences in behavioral choices between the two areas.

**Fig. 3:**
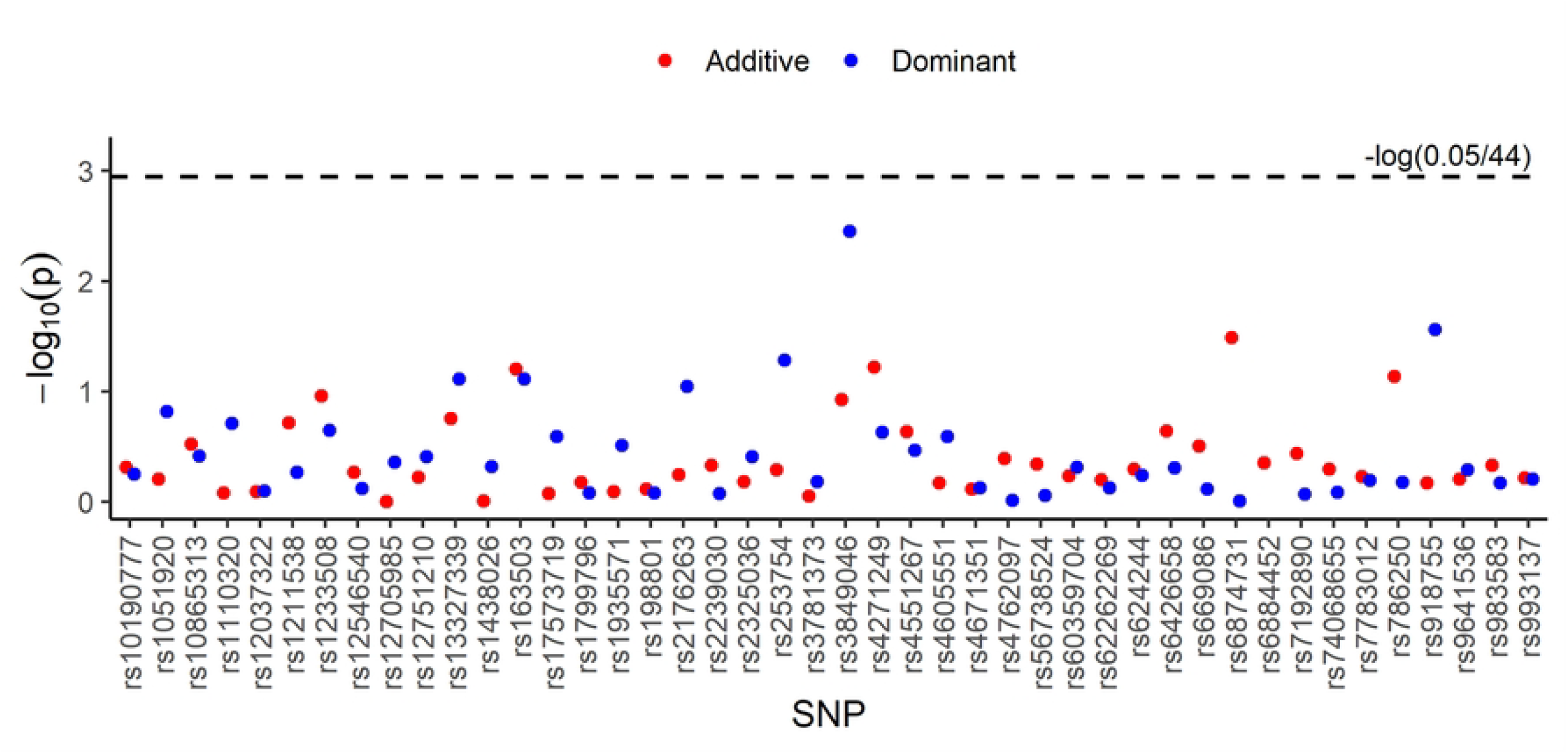
Genetic contribution of 44 candidate SNP on risk tolerance.

Several explanations may account for the lack of support for our hypothesis. First, gene–environment interactions may limit the transferability of GWAS findings across populations. The GWAS by Karlsson Linnér et al. (2019) was conducted primarily in UK and US populations, and genotype–phenotype relationships may differ across environmental and cultural contexts. For example, the lactase persistence allele was strongly associated with stature in ancient populations but not in present-day individuals ^21^. However, a recent study shows that genetic traits influencing personality are robust and highly consistent across geography^22^. Second, risk tolerance is a highly polygenic trait, with individual SNPs generally exerting only small effects. Natural selection may therefore not have had sufficient time to increase the frequency of one or a few variants with effects large enough to be detected in our sample. More generally, the detection of such small genetic effects typically requires very large samples—over one million individuals in the GWAS considered here—which is difficult to reconcile with studies of local adaptation. Third, adaptation to hazardous environments may rely primarily on phenotypic plasticity rather than genetic change. Individuals may adjust their behavior through learning, cultural transmission, developmental processes, or environmentally induced changes in gene regulation, allowing populations to cope with environmental risks without requiring detectable genetic differentiation at the loci examined here. Fourth, the phenotype examined in our study differs from that used in the GWAS by ^7^. Whereas the GWAS relied primarily on self-reported measures of general risk tolerance and risky behaviors, our study employed an incentivized experimental task involving real monetary payoffs. Although both approaches aim to capture individual differences in risk tolerance, they may reflect partly distinct dimensions of the underlying trait. Consequently, genetic variants associated with self-reported risk tolerance may not necessarily predict behavior in incentivized economic decisions.

## Methods

A field study was conducted in the Saint-Louis region in Northern Senegal between March 2018 and March 2020. All experiments were conducted in accordance with relevant guidelines and regulations and informed consent was obtained from all participants. An extension of the first protocol to add SNP genotyping was approved by the Senegalese National Ethics Committee (Comité National d’Ethique en Recherche en Santé) SEN23/47 in 2023.

### Risk-tolerance

Risk tolerance was measured using an incentivized lottery task adapted from Binswanger (1980) as described in ^17^. Participants were invited to choose a card among five. On each card, two amounts were displayed, with an associated color (red or black) and the corresponding amount in coins of XOF 100, in order to have a visual representation. At the end of the experiment, one ball was randomly drawn by a local child and gains were calculated. The cards ranged from completely risk-free (XOF 400 for both balls), to extremely unequal (XOF 0 if Red, XOF 1200 if Black). At each new card, the variance -and the associated risk-is increased, but so is the average amount won.

### MassArray genotyping

DNA was collected on FTA paper and extracted as described in ^17^. Based on the general-risk-tolerance-associated SNPs described in ^7^, we designed three multiplexes with 30, 30 and 23 markers respectively, using the MassARRAY Assay Designer version 4.0.0.2 (Agena Biosciences). All samples were genotyped using the MassARRAY CPM384 system (Agena Bioscience) and the iPLEX Gold chemistry, following ^23^. Allele calling was performed using Typer Analyzer v.5.0.2.137 (Agena Bioscience). We excluded all monomorphic SNPs, loci with weak or ambiguous signals (i.e., displaying more than three clusters of genotypes or unclear cluster delimitation), as well as call rate below 80%.

### Population genetics

SNP loci were tested for conformity with Hardy-Weinberg (HW) equilibrium using the exact probability test ^24^. Deviations from HW equilibrium were measured using the Fis estimator ^25^. SNP loci genotypic differentiation between populations was tested for by calculating an unbiased estimate of the P-value of a log-likelihood (G) based exact test ^26^, a global test over loci was calculated using Fisher’s method. Population differentiation was measured using the Fst estimator ^25^. Calculations were performed using Genepop R package (V. 1.1.7), based on ^27^.

### Statistical analysis

Additive effect is equal to the number (0,1 or 2) of “effect” SNP alleles described in ^7^ and recalled in supplementary data, while the variable dominance effect is a dummy variable equal to 1 when the individual is heterozygote for this SNP or 0 if homozygote. Table S2 shows the beta and p-value of the variable of the following equation:

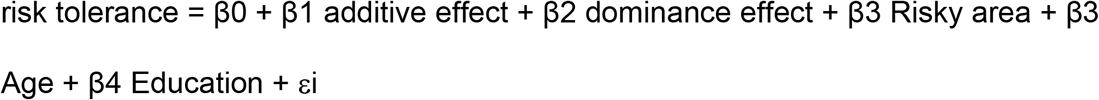

A power analysis was calculated before the genetic analysis was performed. Using the formula described in ^28^, 380 samples allow the detection of an additive effect of β1 = 0.3, which is compatible for a SNP under local adaptation ^29,30^.

## Acknowledgments

The authors also acknowledge the help in data collection from field research assistants. MassArray genotyping was performed at the PGTB (https://doi.org/10.15454/1.5572396583599417E12).

## Author contributions

G-J.C., A.M., B.D., O.S. and M.W. collected the data. G-J.C., C.B., C.M., C.F., A.D, E.G., G.H., M.R. and M.W. analyzed the data. G-J.C., C.M., C.F., M.W. and M.R. wrote the paper. All authors reviewed the manuscript.

## Supporting information

**S1 Table : Differentiation between zone for each SNP**

**S2 Table : GLM correlation between SNPs and risk tolerance**.

